# The nuclear role of methionine adenosyltransferase 2A in immunoglobulin class switch recombination

**DOI:** 10.1101/2025.10.24.684368

**Authors:** Yasuka Ikura, Hiroki Shima, Kanji Furuya, Daisuke Saigusa, Kazuhiko Igarashi

## Abstract

Methionine adenosyltransferase synthesizes S-adenosylmethionine (SAM), the universal methyl donor. Its ubiquitous isozyme MAT2 is present in the nucleus and regulates gene expression through histone methylation. However, the physiological significance of its nuclear enzymatic activity remains unclear. Here, we investigate the role of nuclear MAT2 in antibody class switch recombination (CSR), a crucial process in adaptive immunity. We established murine B-cell lines expressing either wild-type (WT) or catalytically-inactive (D134A) MAT2A subunits, targeted to the nucleus with nuclear localization signal (NLS). Inhibition of nuclear MAT2 enzymatic activity by expressing NLS-MAT2A(D134A) significantly suppressed CSR from IgM to IgA and reduced intracellular SAM levels. Chromatin immunoprecipitation revealed that MAT2A localized to the IgA switch region. Interestingly, while the expression of germline transcripts was enhanced, the expression of activation-induced cytidine deaminase (AID), an essential enzyme for CSR, was downregulated in NLS-MAT2A(D134A) cells. Our findings demonstrate that the enzymatic activity of MAT2 in the nucleus is indispensable for CSR, likely by ensuring proper AID expression subsequent to germline transcription, thus revealing a critical link between nuclear SAM metabolism and adaptive immunity.

## Introduction

S-adenosylmethionine (SAM) is a pivotal metabolite that serves as the primary methyl group donor for the methylation of a wide array of macromolecules, including proteins, lipids, and nucleic acids (Lu, 2000). The synthesis of SAM from methionine and ATP is catalyzed by the enzyme methionine adenosyltransferase (MAT)(Igarashi & Katoh, 2013; Kotb *et al*, 1997). In mammals, two distinct catalytic subunits are expressed from two genes, *MAT1A* and *MAT2A,* generating three isozymes of MAT: MAT1, MAT2, and MAT3. MAT1 and MAT3 are predominantly expressed in the liver and composed of MAT1A subunit, whereas MAT2 is widely distributed in extrahepatic tissues and is a heterooligomer composed of two catalytic subunits, MAT2A, and one regulatory subunit, MAT2B, which is encoded by the *MAT2B* gene (Igarashi & Katoh, 2013; Kotb *et al*., 1997). The MAT2B subunit enhances catalytic activity and stability of MAT2A (Bailey *et al*, 2021; LeGros *et al*, 1997; Markham & Pajares, 2009; Wan *et al*, 2024).

Given the central role SAM in linking one-carbon metabolism to the epigenetic landscape, its intracellular concentration is maintained within a remarkably narrow and optimal range through sophisticated homeostatic mechanisms (Flaherty *et al*, 2025). This tight control is crucial, as both depletion and excess of SAM can be detrimental to cellular health. A key aspect of this regulation occurs at the level of MAT2A itself. It has been well-established that cellular SAM levels govern the splicing and stability of *MAT2A* mRNA; when SAM is abundant, splicing of MAT2A mRNA is reduced and the MAT2A transcript is destabilized and degraded, providing a rapid post-transcriptional feedback loop to prevent over-accumulation of SAM (Shima *et al*, 2017). This intricate control highlights the evolutionary pressure to maintain SAM homeostasis, ensuring metabolic stability while allowing for appropriate epigenetic responses.

Beyond its canonical role in the cytoplasm, including promotion of translation by ribosome by supporting methylation of translation regulators (Alam *et al*, 2022), a sub-pool of MAT2 translocates into the nucleus. Within this compartment, MAT2 is thought to locally synthesize SAM in close proximity to chromatin, thereby providing an efficient source of methyl groups for histone methyltransferases. This nuclear activity directly links the cell’s metabolic state to the epigenetic regulation of gene expression (Katoh *et al*, 2011). Although this process has been implicated in controlling specific gene programs, its broader physiological significance, particularly in dynamic cellular processes that require extensive epigenetic reprogramming, remains largely unexplored.

One such process is the humoral immune response, a cornerstone of adaptive immunity orchestrated by B lymphocytes. Upon activation, B cells can undergo antibody class switch recombination (CSR), a programmed DNA recombination event that diversifies the effector functions of antibodies by replacing the constant region of the immunoglobulin heavy chain (Harriman *et al*, 1993; Lee *et al*, 1998; McHeyzer-Williams & McHeyzer-Williams, 2005). The execution of CSR is dependent on the enzyme Activation-Induced cytidine Deaminase (AID), which initiates the DNA cleavage cascade (Arudchandran *et al*, 2008; Muramatsu *et al*, 2000). CSR is a high-risk process, as it involves the generation of double-strand DNA breaks, which, if improperly repaired, can lead to genomic instability, somatic mutations and oncogenic translocations (Casellas *et al*, 2016; Meng *et al*, 2014; Yu & Lieber, 2019). Consequently, CSR must be exquisitely regulated, both spatially and temporally. The expression of germline transcripts (GLTs), which precedes recombination, peaks within the first few days of an immune response and rapidly declines as B cells commit to CSR. This defines a critical, narrow window during which CSR is licensed to occur (Harriman *et al*., 1993; Lebman *et al*, 1990; Lutzker *et al*, 1988; Stavnezer *et al*, 1988).

Massive metabolic and epigenetic reprogramming are required for B-cell activation, proliferation, and differentiation (Catala-Moll *et al*, 2021; Haniuda *et al*, 2020; Ochiai *et al*, 2024). These massive alterations—all occurring within a short timeframe—undoubtedly places high demands on cellular metabolism, particularly one-carbon metabolism. Indeed, MAT2A expression is strongly induced upon B-cell activation, likely to meet the increased demand for SAM required for extensive histone and DNA methylation changes (Ishii Y et al, submitted). While chemical inhibition of MAT2 has been shown to suppress antibody production (Ishii Y et al, submitted), it has remained unclear whether the critical enzymatic activity of MAT2 is executed in the cytoplasm or the nucleus, and its specific function within the tightly regulated molecular cascade of CSR has not been elucidated.

Given that CSR is a temporally restricted and tightly controlled process dependent on epigenetic modifications, we hypothesized that nuclear SAM synthesis would constitute a critical regulatory node during this specific, early window of B-cell activation by providing SAM locally for efficient epigenetic modifications. In this study, we aimed to dissect the specific biological role of the enzymatic activity of nuclear MAT2 in humoral immunity, using CSR as a model system, by delivering catalytically inactive MAT2A in the nucleus. Our results provide direct evidence that nuclear SAM synthesis by MAT2 is indispensable for CSR and required for the proper induction of AID. Furthermore, we uncover that a lack of nuclear SAM synthesis induces the feedback mechanism regulating MAT2A mRNA.

## Results

### Establishment of nuclear expression systems for wild-type and mutant MAT2A in CH12F3-2A cells

The catalytic site of MAT2 is formed at the interface of MAT2A dimer. (Shafqat *et al*, 2013). Therefore, by mutating residues critical for the catalysis, one may generate a dominant negative mutation which would interfere the catalytic activity of wild-type subunit as well. To investigate the role of MAT2’s enzymatic activity within the nucleus, we established murine B-cell lymphoma CH12F3-2A cell lines engineered to express Flag-HA-tagged wild-type (WT) or catalytically inactive (D134A) MAT2A, each fused to a nuclear localization signal (eNLS: epitope-tagged NLS). Western blot analysis of the stably expressing cell lines confirmed the expression of the exogenous eNLS-MAT2A(WT) and eNLS-MAT2A(D134A) proteins, which were distinguishable by their higher molecular weight from the endogenous MAT2A (**Fig. 1A**). The amounts of exogenous MAT2A were less than endogenous MAT2A. Immunofluorescence staining demonstrated their successful targeting of the eNLS-MAT2A proteins to the nucleus, in contrast to the predominantly cytoplasmic localization of exogenous MAT2A proteins lacking an NLS (**Fig. 1B**). This specific nuclear accumulation validated our strategy to uncouple the nuclear and cytoplasmic functions of the enzyme, thereby creating a new system to assess the role of nuclear MAT2.

**Figure 1.**
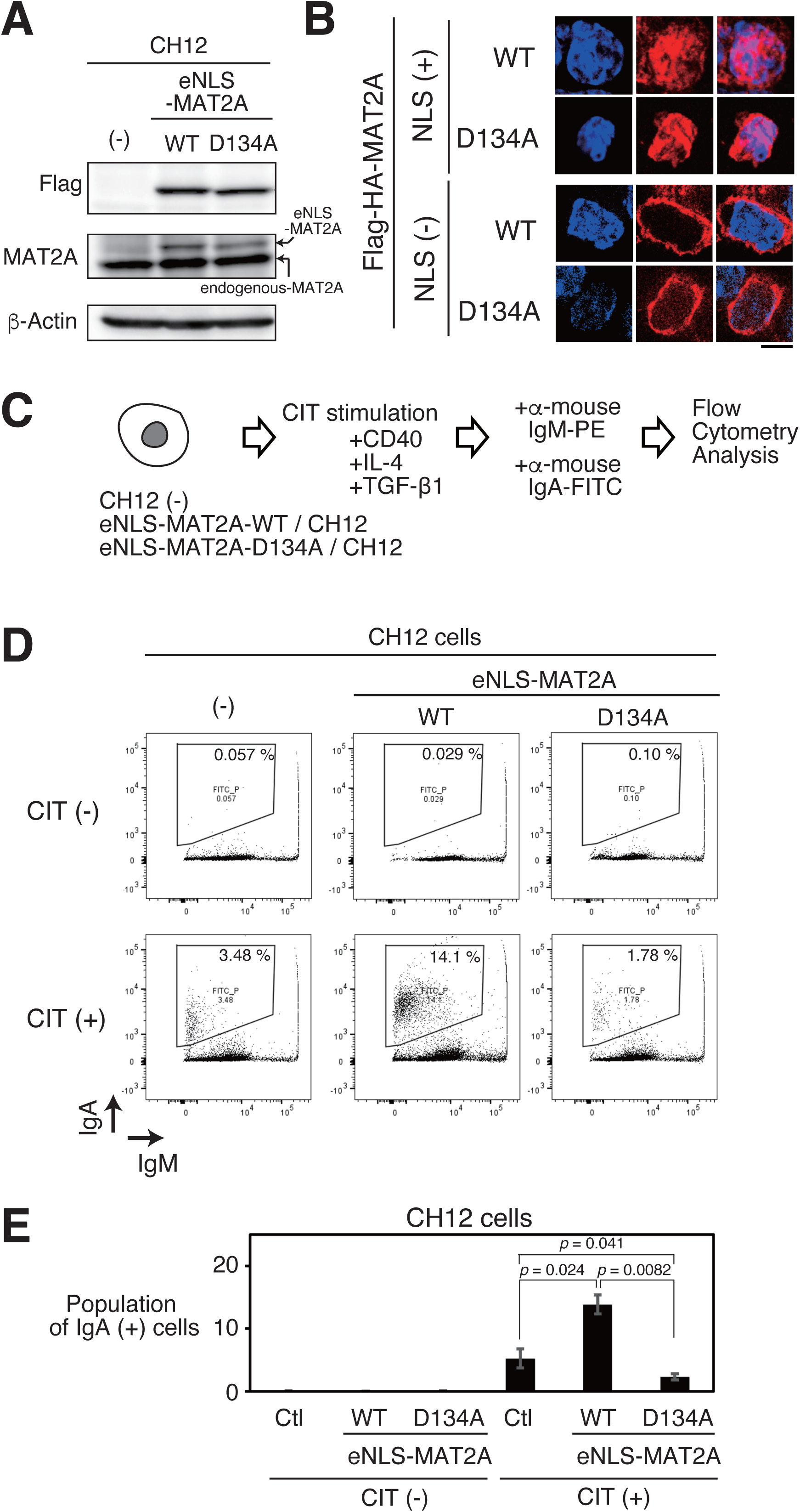
Nuclear expression of MAT2A mutant inhibits CIT-induced class switch recombination from IgM to IgA in CH12F3-2A cells. **A.** Expression of Flag-HA-tagged NLS-MAT2A(WT) (eNLS-MAT2A(WT)) and NLS-MAT2A(D134A) (eNLS-MAT2A(D134A)) proteins in CH12F3-2A cells was confirmed by western blotting using the indicated antibodies. **B.** Nuclear localization of Flag-HA-tagged NLS-MAT2A(WT) and NLS-MAT2A(D134A) proteins was analyzed by immunofluorescence staining using an anti-Flag antibody. NLS (+): cells expressing MAT2A with an added NLS; NLS (–): cells expressing MAT2A without NLS; WT: cells expressing wild-type MAT2A; D134A: cells expressing catalytically inactive MAT2A(D134A) mutant. Red: FLAG; Blue: DNA (DAPI). Bar, 20 µm. **C.** Schematic representation of the experimental workflow for CD40, IL-4, and TGF-β1 (CIT)-induced class switch recombination from IgM to IgA in CH12F3-2A cells. Wild-type CH12F3-2A cells, eNLS-MAT2A(WT)/CH12F3-2A cells, and eNLS-MAT2A(D134A)/CH12F3-2A cells were stimulated with CIT, and IgA expression was analyzed by flow cytometry at 48 and 72 hours after stimulation. **D.** Scatter plots of flow cytometry analysis showing IgM (PE) signal on the x-axis and IgA (FITC) signal on the y-axis. The solid-line box indicates the gate for IgA-positive cells, and the numbers in each panel represent the percentage of IgA-positive cells under each condition. **E.** Bar graphs showing time-dependent changes in the proportion of IgA-positive cells in each cell line. Data represent means of three biological replicates. Error bars indicate SD. *p*-values were calculated using Student’s *t*-test.

### Nuclear expression of a catalytically inactive MAT2A mutant inhibits CIT-induced class switch recombination

We next examined the functional consequence of inhibiting nuclear MAT2A activity on antibody class switch recombination (CSR). As shown in the experimental schematic (**Fig. 1C**), CH12F3-2A cells were stimulated with a cocktail of CD40 ligand, IL-4, and TGF-β1 (abbreviated as CIT) to induce CSR from IgM to IgA (Lee *et al*, 2023; Nakamura *et al*, 1996). After 72 hours of stimulation, flow cytometry analysis revealed that roughly 5% of the control cells without any exogenous MAT2A became IgA-positive. Overexpression of eNLS-MAT2A(WT) increased the number of cells with CSR, up to 15%. Expression of the eNLS-MAT2A(D134A) mutant resulted in a profound near-complete inhibition of the switch to IgA, down to less than 2% (**Fig. 1D, E**). These findings strongly indicates that nuclear SAM is limiting for CSR in CH12F3-2A cells and that the enzymatic activity of MAT2A within the nucleus is essential for the successful execution of CSR.

### Nuclear MAT2A activity is critical for maintaining SAM homeostasis during class switch recombination

To understand the biochemical basis for the CSR defect, we analyzed SAM-related metabolites following CIT stimulation using mass spectrometery (**Fig. 2A**). In wild-type CH12F3-2A cells, SAM levels decreased over time regardless of CIT stimulation. Conversely, methionine (Met), a precursor for SAM synthesis, and S-adenosylhomocysteine (SAH), a metabolite derived from SAM, showed a significant increase only after 72 hours under CIT stimulation (**Fig, 2B**), which suggest that SAM metabolism shifts from “synthesis” to “consumption” during antibody gene class switch recombination in wild-type CH12F3-2A cells.

**Figure 2.**
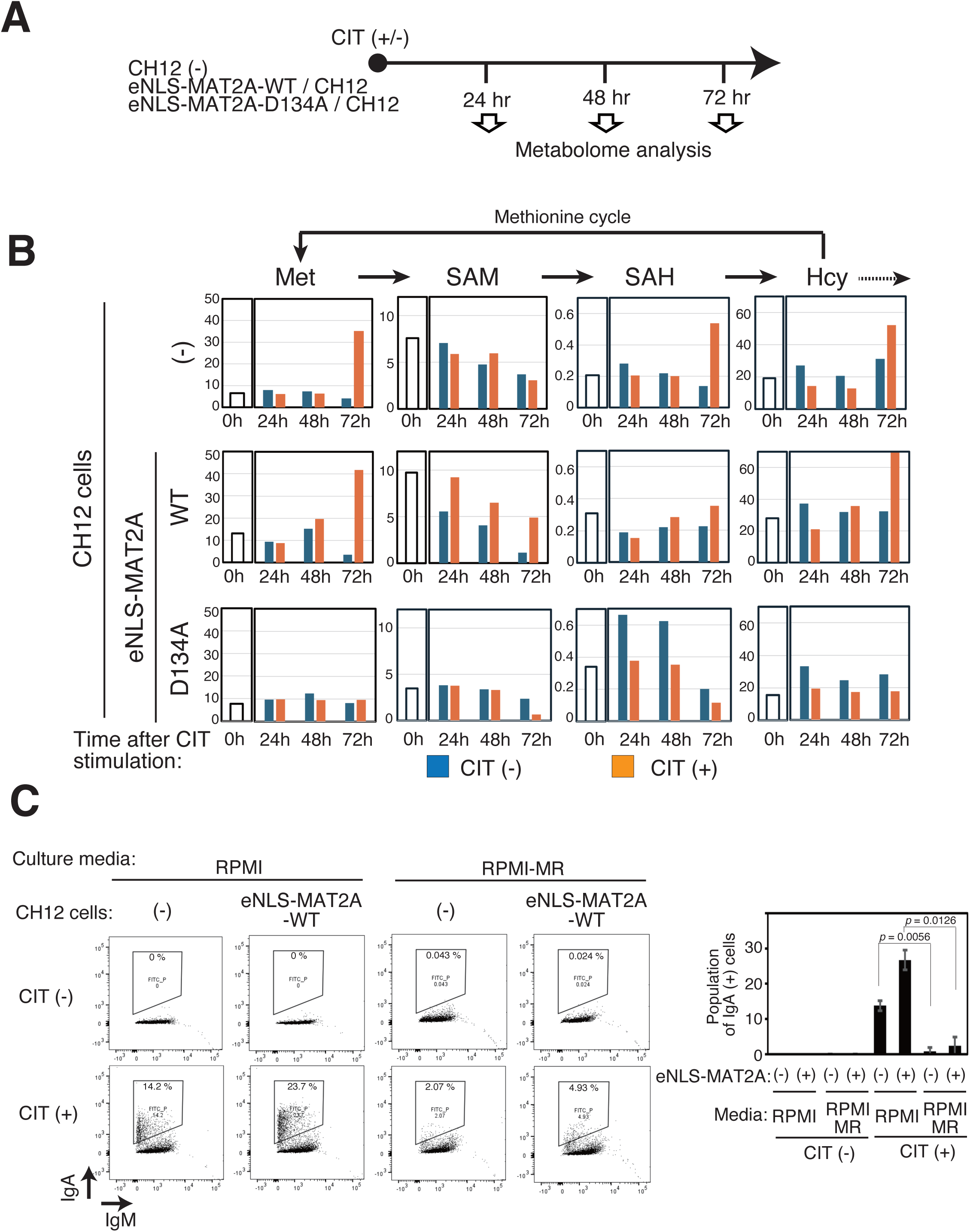
Nuclear expression of MAT2A mutant alters CIT-induced class switch recombination via methionine metabolism reprogramming in CH12F3-2A cells. **A.** Schematic representation of the experimental workflow for analyzing SAM-related metabolites following CD40, IL-4, and TGF-β1 (CIT)-induced class switch recombination in wild-type CH12F3-2A cells, eNLS-MAT2A(WT)/CH12F3-2A cells, and eNLS-MAT2A(D134A)/CH12F3-2A cells. **B.** Time-course analysis of SAM-related metabolites in each cell line after CIT stimulation. Methionine (Met), S-adenosylmethionine (SAM), S-adenosylhomocysteine (SAH), and homocysteine (Hcy) are shown. **C.** Class switch recombination under methionine-restricted conditions. Wild-type CH12F3-2A cells, eNLS-MAT2A(WT)/CH12F3-2A cells, and eNLS-MAT2A(D134A)/CH12F3-2A cells were cultured in methionine-restricted medium and subjected to CIT stimulation. Scatter plots of flow-cytometry analysis show IgM (PE) signal on the x-axis and IgA (FITC) signal on the y-axis. The solid-line box indicates the gate for IgA-positive cells, and the numbers in each panel represent the percentage of IgA-positive cells under each condition. Bar graphs show the quantified proportion of IgA-positive cells in each condition. Under methionine-restricted conditions, CIT-induced IgA class switch recombination was markedly suppressed. Data represent means of three biological replicates. Error bars indicate SD. *p*-values were calculated using Student’s *t*-test.

Furthermore, eNLS-MAT2A(WT)/CH12 cells exhibited higher SAM levels than wild-type CH12F3-2A cells specifically upon CIT stimulation. In contrast, eNLS-MAT2A(D134A)/CH12 cells displayed lower SAM levels compared to both wild-type CH12F3-2A cells and eNLS-MAT2A(WT)/CH12 cells (**Fig, 2B**). Collectively, these results indicate a crucial role for MAT2A-dependent nuclear SAM synthesis in antibody gene class switch recombination. The indispensable nature of SAM synthesis was further validated by showing that CSR was markedly suppressed when wild-type CH12F3-2A cells were cultured in a methionine-restricted medium, a condition which phenocopies the metabolic state of the NLS-MAT2A(D134A) mutant (**Fig. 2C**).

### Inhibition of nuclear SAM synthesis induces a transcriptional feedback loop that upregulates endogenous MAT2A

The severe depletion of SAM in eNLS-MAT2A(D134A) cells led us to investigate whether the feedback regulation of MAT2A mRNA was operative. We discovered that the expression of endogenous MAT2A protein was significantly and progressively upregulated in eNLS-MAT2A(D134A) cells whereas it was reduced in eNLS-MAT2A(WT) cells (**Fig. 3A**). These upregulation and down regulation were driven at the transcriptional level, as real-time RT-PCR revealed corresponding alterations of endogenous Mat2a mRNA (**Fig. 3B**). These alterations were consistent with the known post-transcriptional regulation of mRNA stability to maintain SAM levels (Pendleton *et al*, 2017; Shima *et al*., 2017). Considering the persistent SAM depletion seen in **Fig. 3B**, they also suggested that the function of the increased endogenous MAT2A was effectively interfered by eNLS-MAT2A(D134A).

**Figure 3.**
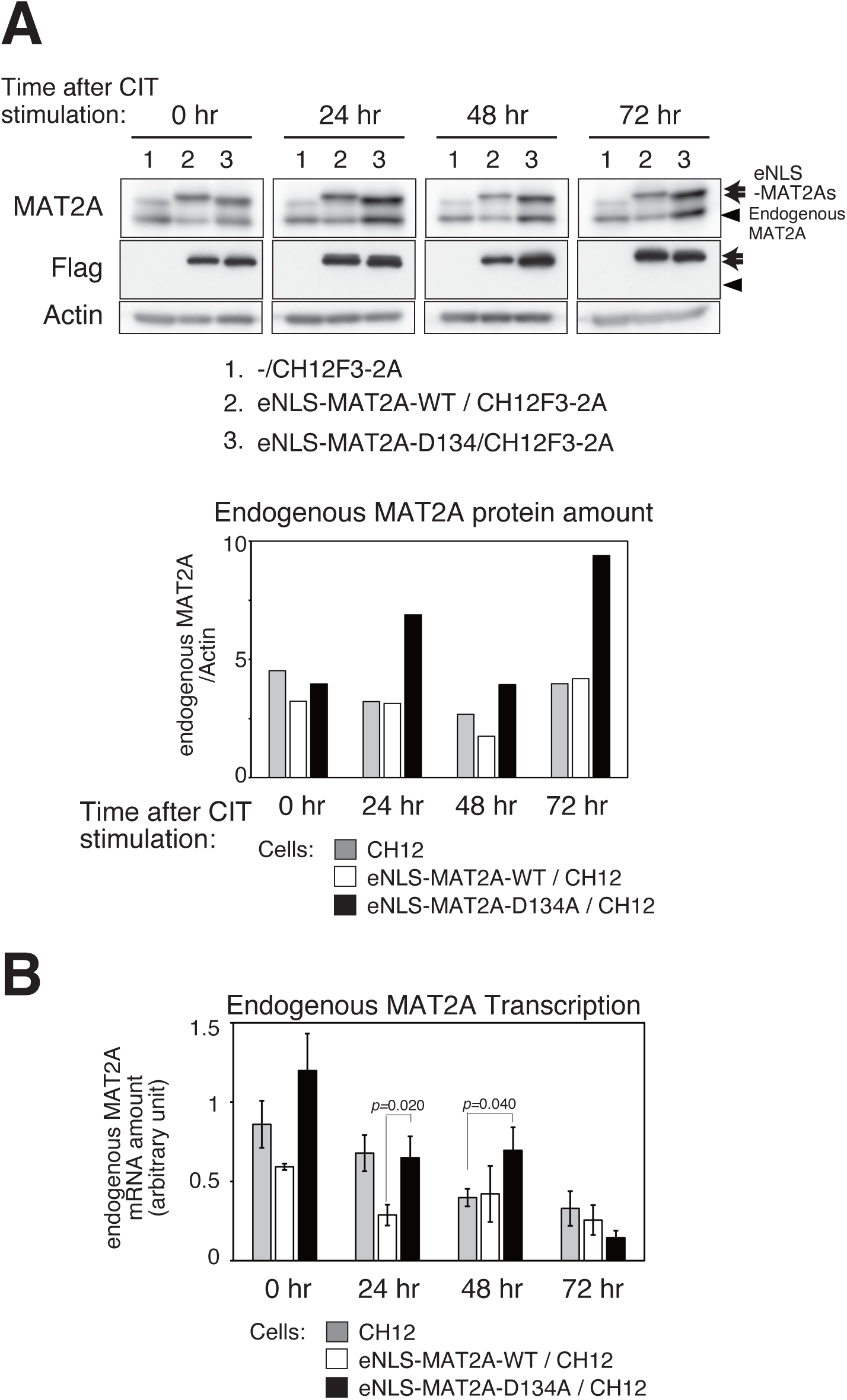
Inhibition of nuclear MAT2A-dependent SAM synthesis induces endogenous MAT2A expression. **A.** Expression of endogenous MAT2A protein (black arrowhead) and NLS-MAT2A(WT)/(D134A) proteins (arrows) in CH12F3-2A, eNLS-MAT2A(WT)/CH12F3-2A, and eNLS-MAT2A(D134A)/CH12F3-2A cells collected at the indicated time points after CIT stimulation was analyzed by western blotting. The bar graphs show the quantification of endogenous MAT2A protein levels normalized to actin. **B.** Expression of endogenous MAT2A mRNA after CIT stimulation was quantified by real-time RT–PCR using cDNA synthesized from total RNA (2 µg per sample) under the indicated conditions. The relative expression levels were normalized and expressed in arbitrary units (see *Methods* for details). Data represent means of three biological replicates. Error bars indicate SD. *p*-values were calculated using Student’s *t*-test.

### Disrupted nuclear SAM homeostasis leads to aberrant H3K36 methylation at the IgA switch region

To dissect the epigenetic consequences of this disrupted metabolic state, we performed chromatin immunoprecipitation (ChIP) analysis. Analysis of wild-type CH12F3-2A cells revealed that MAT2A preferentially localized to the IgA switch (Iα) region, a target for class switching, even before CIT stimulation, compared to the IgM (Iµ) region **(Fig. 4A, B)**. Intriguingly, MAT2 exhibited a stronger association with chromatin in cells expressing eNLS-MAT2A(D134A) than in those expressing the wild-type counterpart, eNLS-MAT2A(WT), possibly reflecting a lack of enzymatic turnover that ‘traps’ the inactive protein on its chromatin targets **(Fig. 4C).** We next measured key histone methylation. While H3K4me3 and H3K79me3 levels were modestly increased in eNLS-MAT2A(WT) cells, this increase was not observed eNLS-MAT2A(D134A) cells **(Fig. 4D)**. In contrast, we observed a significant increase in H3K36 trimethylation (H3K36me3) specifically at the downstream of Iµ and at Iα region in eNLS-MAT2A(D134A) cells **(Fig. 4D)**. These results suggested that the failure of SAM homeostasis did not cause a global loss of methylation, but rather led to a specific and aberrant alteration of the histone modification at the target site of recombination.

**Figure 4.**
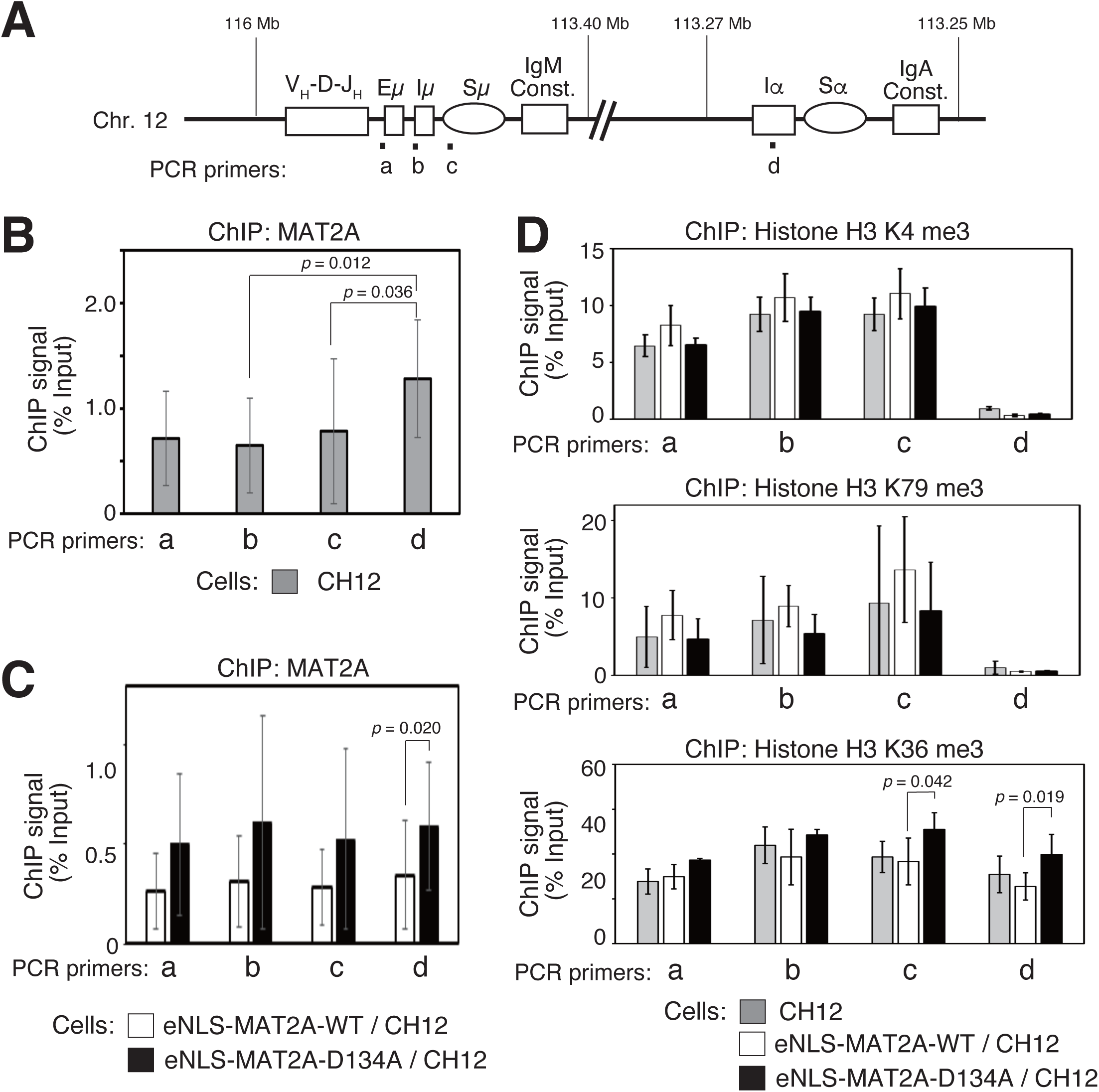
Chromatin immunoprecipitation analysis reveals MAT2A localization and distinct H3K36 methylation patterns at the I-alpha locus. **A.** Schematic representation of the immunoglobulin heavy chain (Igh) gene locus showing the approximate locations of primers used for ChIP analysis. **B.** ChIP analysis of endogenous MAT2A in wild-type CH12F3-2A cells. Endogenous MAT2A was found to localize at the IgA switch (Ia) region. CIT stimulation was not applied, indicating that endogenous MAT2A is already associated with the Ia region prior to CIT induction. **C.** ChIP analysis using an anti-MAT2A antibody in eNLS-MAT2A(WT)/CH12F3-2A and eNLS-MAT2A(D134A)/CH12F3-2A cells under unstimulated conditions. The NLS-MAT2A(D134A) mutant protein exhibited stronger chromatin association compared with NLS-MAT2A(WT). **D.** ChIP-qPCR analyses of CH12F3-2A, eNLS-MAT2A(WT)/CH12F3-2A, and eNLS-MAT2A(D134A)/CH12F3-2A cells using antibodies against histone H3K4me3, H3K79me3, and H3K36me3. B-D. Data represent means of three biological replicates. Error bars indicate SD. *p-* values were calculated using Student’s *t*-test.

### Nuclear SAM synthesis is critical for coupling germline transcription to AID expression

We next examined the expression of α germline transcripts (αGLT) (**Fig. 5A**). Both of the long and short forms of αGLT was transiently increased in the control CH12F3-2A cells. These GLTs were increased in both eNLS-MAT2A(WT) and eNLS-MAT2A(D134A) cells, patterns became different. While the peak level was blunted in eNLS-MAT2A(WT) cells, αGLTs were significantly upregulated and sustained in eNLS-MAT2A(D134A) cells upon CIT stimulation (**Fig. 5B**). This upregulation was specific to αGLTs, as µGLT (which, as previously reported, does not fluctuate with CIT stimulation) and control *Actin* mRNA showed no such changes **(Fig. 5C)**. This presented a central and revealing paradox: despite robust, even enhanced, expression of the transcripts that initiate CSR by opening chromatin, the process itself was completely blocked in eNLS-MAT2A(D134A) cells. This apparent uncoupling of events led us to examine the expression of the master enzyme of CSR, AID. Transcription of AID, encoded by the *Aicda* gene, was reduced in eNLS-MAT2A(WT)/CH12 cells compared to wild-type CH12F3-2A cells, and this induction was further diminished in eNLS-MAT2A(D134A)/CH12 cells **(Fig. 5C)**. Moreover, at the protein level, AID expression was induced in eNLS-MAT2A(WT)/CH12 cells over 72 hours following CIT stimulation, whereas this induction was suppressed in eNLS-MAT2A(D134A)/CH12 cells **(Fig. 5D)**.

**Figure 5.**
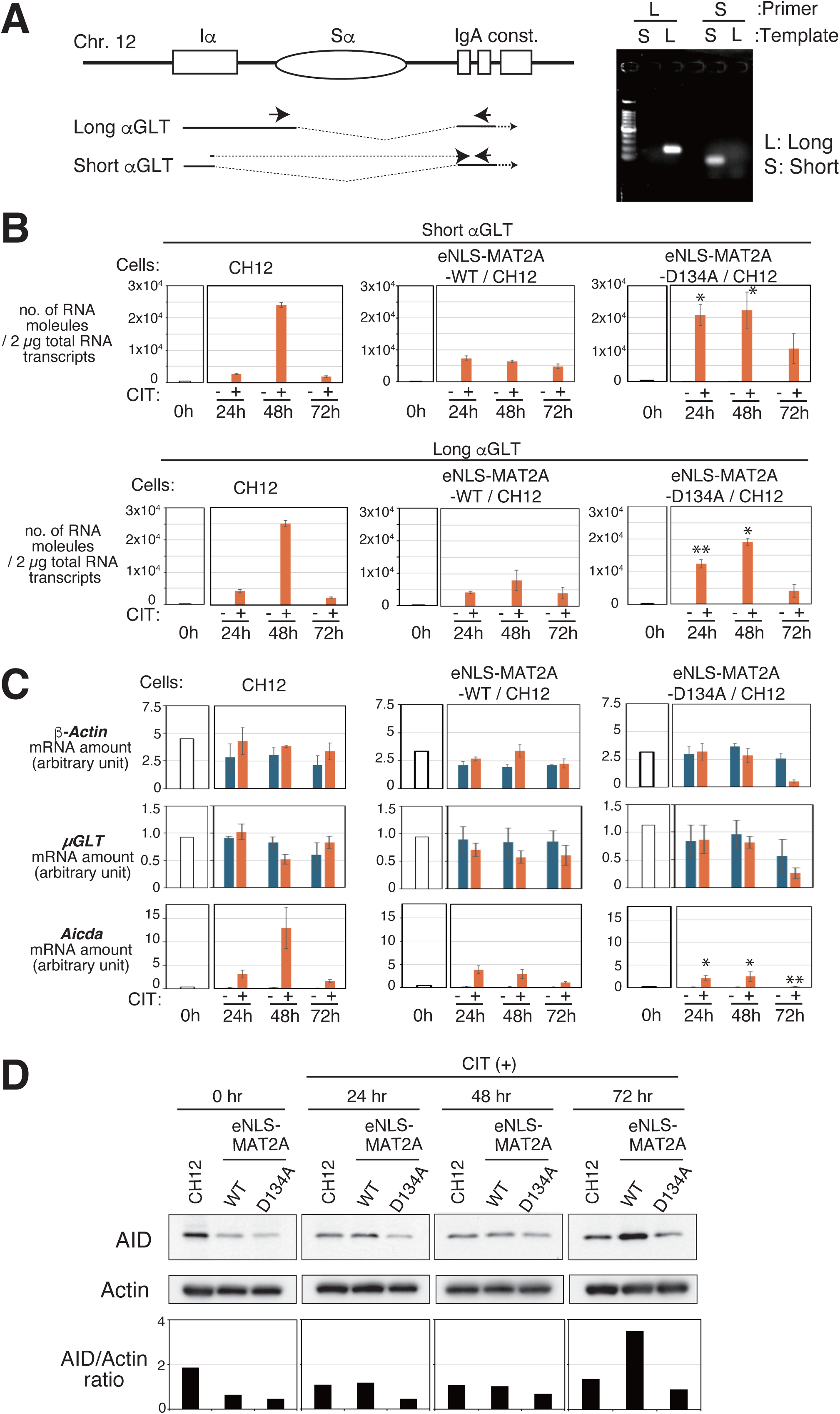
Nuclear SAM synthesis is critical for IgA class switch recombination. A. Schematic representation showing the primer sets used to detect the *αGLT* short and long transcripts. Each primer pair specifically amplifies either the short or long *αGLT* isoform. B. Time-course quantitative PCR (qPCR) analysis of short (upper row) and long (lower row) *αGLT* mRNA expression in CH12F3-2A cells and CH12F3-2A cells expressing eNLS-MAT2A(WT) or eNLS-MAT2A(D134A). The y-axis indicates the number of *αGLT* transcripts per µg of total RNA (see *Methods* for details). Upon CIT stimulation, expression of the short *αGLT* transcript was induced earlier than that of the long *αGLT*, and this induction was more pronounced in eNLS-MAT2A(D134A)–expressing cells. C. Quantitative PCR analysis of *β-Actin*, μGLT, and *Aicda* gene expression in CH12F3-2A cells and CH12F3-2A cells expressing eNLS-MAT2A(WT) or eNLS-MAT2A(D134A), using cDNA synthesized from total RNA (2 µg per sample) under the indicated conditions. Expression levels were normalized relative to the control sample and are presented in arbitrary units (see *Methods* for details). B-C. Data represent means of three biological replicates. Error bars indicate SD. *p*-values were calculated using Student’s *t*-test. * indicates *P* < 0.05; ** indicates *P* < 0.01. D. Western blot analysis of AID (*Aicda* gene product). Upper panel: Immunoblot of AID using the indicated antibody. Lower panel: Quantification of AID protein levels normalized to actin.

## Discussion

In this study, we have delineated an essential role for the nuclear enzymatic activity of MAT2 in antibody class switch recombination. Our findings point to the importance of the nuclear synthesis of SAM and its feedback network, which rigorously control intracellular SAM levels for the successful execution of CSR in B cells. We showed that inhibiting the enzymatic function of MAT2 specifically within the nucleus leads to a profound CSR defect in the model CH12 B cells. The biochemical underpinning of this defect was a failure to maintain cellular SAM homeostasis during activation of CH12 cells. Our experimental results highlight that the nuclear compartment is not merely a passive recipient of cytoplasmic SAM but is a site of active and essential SAM synthesis.

While the eNLS-MAT2A(D134A) cells showed reductions in the amount of SAM compared to the control or eNLS-MAT2A(WT) cells, both MAT2A mRNA and protein were increased. These results indicate that the cell actively senses a deficit in nuclear SAM synthesis and responds by upregulating endogenous Mat2a mRNA depending on the known post-transcriptional control via mRNA stability or splicing. The failure of the cells to restore SAM amount shows that eNLS-MAT2A(D134A) effectively inhibited the enzymatic activity of endogenous nuclear MAT2. The profound CSR defect in these cells is therefore best understood not as a simple consequence of SAM deficiency, but as a state of feedback control failure. Our results suggest that, when nuclear SAM deficiency is unresolved, nuclear events like CSR program become defective. eNLS-MAT2A(WT) moved cells toward the opposite direction along the SAM amount, where CSR was increased. Taken together, these results strongly suggests that SAM is a limiting factor for CSR in CH12 cells and that metabolic status of SAM serves as a crucial gatekeeper of CSR in B cells.

How did MAT2A(D134A) and resulting SAM deficiency cause the CSR failure? Or how did NLS-MAT2A(D134A) cause CSR increase? The concurrent hyper-induction of H3K36me3 and αGLT in MAT2A(D134A)-expressing cells may be explained by the reduction of CSR in these cells. A reduction of DNA breakage within the switch regions would increase these measurements. Consistent with this idea, αGLT was decreased in MAT2A(WT)-expressing cells. The specific enhancement of H3K36me3 in a state of overall SAM reduction is counterintuitive, but it may reflect a dysfunctional metabolic state where the normally local production of SAM is lost, leading to aberrant activity of specific histone methyltransferases like MMSET (Pei *et al*, 2013). This hypothesis is strongly supported by previous reports linking MMSET activity directly to αGLT transcription (Pei *et al*., 2013), which aligns perfectly with our observation of enhanced αGLT expression. This finding challenges a simple model where GLT expression is the sole rate-limiting step for CSR, and instead points to a more complex regulatory mechanism in which SAM-dependent critical events are placed downstream of GLT-mediated chromatin opening. This could be the induction of AID expression, which was reduced in NLS-MAT2A(D134A) cells. Considering that CSR is a temporally restricted event occurring primarily in a narrow window before or shortly after GC entry, it is logical that its master switch, AID, would be regulated by a mechanism that senses the cell’s overall metabolic and epigenetic readiness. Considering that AID was also reduced in NLS-MAT2A(WT) cells, its expression is regulated in a biphasic manner in response to nuclear SAM. In addition to AID, there may be additional SAM-sensitive event(s) downstream of GLT expression. The MAT2A-SAM feedback loop appears to be a central component of this ‘readiness’ checkpoint. The failure of CSR is not simply a insufficiency of SAM, but a failure of the sophisticated regulatory system that ensures SAM is supplied in a controlled and timely manner for multiple transmethylation reactions.

In conclusion, our work suggests that nuclear synthesis of SAM enables the precise epigenetic control required for the complex, high-risk DNA rearrangements that define vertebrate adaptive immunity. We propose that nuclear SAM homeostasis tunes the probability of CSR in B cells and hence successful execution of the adaptive immune response, realizing metabolic control of immune response.

## Methods

### Cell culture and class switch induction

CH12F3-2A cells were obtained from the RIKEN Cell Bank (Cell Engineering Division, RIKEN; Kato H. *et al*., 1997). Cells were cultured in RPMI-1640 medium with L-glutamine and phenol red, supplemented with 5% NCTC-109, 20 mM HEPES, 10% fetal bovine serum, and 0.055 mM 2-mercaptoethanol, at 37 °C in a humidified atmosphere containing 5% CO₂.

For the class switch induction, wild-type CH12F3-2A cells were cultured in RPMI1640 medium supplemented with CD40 (final concentration 2 μg/mL; Invitrogen, CD40 Monoclonal Antibody [1C10], eBioscience™), IL-4 (final concentration 10 ng/mL; BD Pharmingen™, Recombinant Mouse IL-4), and TGF-β1 (final concentration 1 ng/mL; R&D Systems, Recombinant Mouse TGF-beta 1, 7666-MB-005)(Lee *et al*., 2023).

### Construction of NLS-MAT2A expression cells

pOZ-N-NLS-MAT2A(WT) and pOZ-N-NLS-MAT2A(D134A) constructs were generated using the pOZ-N retroviral vector(Ikura *et al*, 2000; Nakatani & Ogryzko, 2003), which contains N-terminal Flag and HA tags. The SV40 NLS sequence (PPKKKRKV) was inserted immediately downstream of the FLAG and HA tags and upstream of the XhoI site, followed by the first methionine codon of MAT2A. The resulting retroviral vectors were introduced into CH12F3-2A cells, and stable cell lines expressing eNLS-MAT2A(WT) or eNLS-MAT2A(D134A) were established as previously described.

### Mass Spectrometry

Cells were collected and frozen in -80℃. The sample was deproteinized by adding the methanol solution (100 μl) containing the internal standards; GSH-^13^C_2_,^15^N and SAM-^13^C_5_,^15^N (1 μg/ml, each). After mixing for 30 sec, sonicating for 5 min and centrifuging at 16,400 × *g* for 10 min at 4℃, the supernatant was transferred to the vial for the ultra-high-performance liquid chromatography triple quadrupole mass spectrometry (UHPLC-MS/MS). The UHPLC-MS/MS was performed with the I-class UPLC system conducted to Xevo TQ-XS (Waters corp. Cheshire, UK). The detailed analytical condition and data processing were previously described (Nishizawa *et al*, 2020).

### Immunoblot analysis and antibodies

The following primary antibodies were used for immunoblotting: anti-MAT2A (Proteintech, 55309-1-AP; 1:1000 dilution), anti-Flag (Sigma, A8592; 1:4000 dilution), anti-β-Actin (GeneTex, GTX109639; 1:1000 dilution), and anti-AID (Cell Signaling Technology, #4975; 1:1000 dilution).

The preparation of cell lysates for immunoblot analysis was performed using RIPA buffer (10 mM Tris-HCl [pH 7.4], 150 mM NaCl, 5 mM EDTA, 1% Triton X-100, 1% sodium deoxycholate, 0.1% sodium dodecyl sulfate, and 0.2 mM PMSF) as the extraction buffer.

### Fluorescence Immunostaining

Fluorescence immunostaining of suspension cells was performed following the method established by Miyazawa *et al*. (2025)(Miyazawa *et al*, 2025), with slight modifications for CH12F3-2A cells.

Briefly, 3.5 × 10⁵ cells were transferred to a 1.5-ml tube and centrifuged at 100 × *g* for 1 min at 25 °C (KUBOTA 3740). The cells were washed twice with PBS and applied onto a PVDF membrane–based centrifugal filter unit (Ultrafree-MC, 0.65 μm pore size, Millipore). After washing with PBS, cells were fixed in 4% paraformaldehyde for 30 min at room temperature, washed twice with PBS, and blocked with 1% BSA/PBS for 30 min.

Primary antibody incubation was performed for 1 h at room temperature using anti-FLAG (Sigma-Aldrich, Monoclonal ANTI-FLAG M2, F3165; 1:1000 dilution). After two PBS washes, cells were incubated with a Cy3-conjugated donkey anti-mouse IgG (H+L) secondary antibody (Jackson ImmunoResearch, 715-165-151; 1:2000 dilution) for 30 min at room temperature, followed by PBS washes. Nuclei were stained with DAPI (Roche, 10236276001) for 1 min.

The PVDF membrane containing the stained cells was mounted on a glass slide with 90% glycerol and covered with a coverslip. Images were acquired using a Keyence BZ-X710 all-in-one fluorescence microscope.

### Chromatin Immuno-Precipitation (ChIP) analysis

CH12F3-2A cells expressing eNLS-MAT2A (WT), eNLS-MAT2A (D134A), or wild-type CH12F3-2A cells were cultured in 10 cm dishes until reaching a density of 1 × 10⁶ cells/mL. Cells were collected by centrifugation at 1500 rpm for 5 min at 4 °C (KUBOTA 5910, swing rotor). Cell pellets were resuspended in 5 mL of ice-cold 1% paraformaldehyde (PFA) in RPMI1640 for fixation and incubated on ice for 10 min. Fixation was quenched by resuspension in 125 mM glycine, followed by centrifugation at 1500 rpm for 5 min at 4 °C. Pellets were washed with FACS buffer (FACS buffer, 1×PBS(-), 2% FBS, 0.05% NaN_3,_ 200 nM PMSF), mixed on a MACSmix™ Tube Rotator at 4 °C for 5 min, and centrifuged. Cells were lysed in SDS lysis buffer (50 mM Tris-HCl [pH 8.0], 10 mM EDTA, 1%SDS, 200 nM PMSF) and incubated for 10 min at room temperature. Chromatin was sonicated using a Bioruptor (Cosmo Bio) with five cycles of 10 s sonication followed by 1 min incubation on ice. Lysates were clarified by centrifugation at 15,000 rpm for 10 min at 8 °C (Himac CT15RE). Supernatants were diluted with ChIP dilution buffer (50 mM Tris-HCl [pH 8.0], 167 mM NaCl, 1.1% Triton X-100, 0.11% NaDoc, 200 nM PMSF), precleared with normal rabbit IgG (Cell Signaling Technology, #2729) or normal mouse IgG (Santa Cruz, sc-2025), and incubated with Protein G agarose beads (Thermo Scientific, 20398) pre-washed in 3.125 mg/mL BSA/ChIP dilution buffer for 2 h at 4 °C. Samples after this step were designated as “Input”.

For Antibody conjugation, Pierce™ Protein A/G magnetic beads (Thermo Scientific) were washed with PBN buffer (10× PBS(-), 10% BSA, 1% NaN1×PBS(-), 0.5%BSA, 0.05%NaN_3_) and incubated overnight at 4 °C with the following antibodies: anti-MAT2A (Proteintech, 55309-1-AP), anti-Histone H3K4me3 (Abcam, ab8580), anti-Histone H3K36me3 (Abcam, ab9050), and anti-Histone H3K79me3 (Abcam, ab2621). The antibody-conjugated beads were washed with PBN buffer and 1× RIPA buffer/150 mM NaCl before mixing with Input samples. The mixture was incubated overnight at 4 °C on a MACSmix™ Tube Rotator. Beads were sequentially washed with 1× RIPA buffer/150 mM NaCl, 1× RIPA buffer/500 mM NaCl, LiCl wash buffer (10 mM Tris-HCl [pH 8.0], 1 mM EDTA, 250 mM LiCl, 0.5% NP-40, 0.5% NaDoc), and twice with TE buffer. DNA-protein complexes were eluted with ChIP elution buffer (10 mM Tris-HCl[pH 8.0], 5 mM EDTA, 300 mM NaCl, 0.5% SDS) and incubated overnight at 65 °C to reverse cross-links. The eluates were treated with RNase A at 37 °C for 30 min, followed by Proteinase K (FUJIFILM Wako, 161-28701) at 55 °C for 2.5 h. IP samples were clarified by centrifugation at 10,000 rpm for 10 s at 8 °C, and supernatants were collected. Finally, DNA was purified using the ChIP DNA Clean & Concentrator kit (Zymo Research, D5205) according to the manufacturer’s instructions.

Purified DNA was analyzed by real-time PCR using a LightCycler 96 system (Roche) and the FastStart SYBR Green Master SYBR Green I (Roche) according to the manufacturer’s instructions. The primers used in ChIOP analysis are listed in Table S1.

### RT-qPCR analysis

Total RNA was prepared using an RNeasy Plus Mini Kit (Quiagen), according to the manufacturer’s instructions. RNA concentration and purity were assessed using a NanoDrop-1000 spectrophotometer (Thermo Fisher Scientific). The extracted RNA was diluted with nuclease-free water to a final concentration of 200 ng/µL (2000 ng in 10 µL). cDNA was synthesized from the diluted RNA using the High-Capacity cDNA Reverse Transcription Kit (Thermo Fisher Scientific, Cat. No. 4374966) following the manufacturer’s protocol. The FastStart SYBR Green Master SYBR Green I (Roche) for cDNA was used for quantitative PCR by LightCycler96 (Roche). The relative expression was normalized by β2-microglobulin (β2 m). Primers used in this study are shown in Table S1.

### Identification and quantitative analysis of αGLT isoforms

Total RNA was isolated from CH12F3-2A cells at 24 h and 72 h after class switch induction with CD40, IL-4, and TGF-β1, and cDNA libraries were synthesized. αGLT transcripts were amplified by nested PCR using outer primers (GLT-Forward and GLT-Reverse) followed by inner primers (GLT-Forward2 and GLT-Reverse2). PCR was performed with PrimeStar Max (Takara, R045A) under the following conditions: 95 °C for 2 min, followed by 30 cycles of 98 °C for 10 s and 68 °C for 4 min. PCR products were separated by agarose gel electrophoresis, visualized with Sybr Green, and yielded two distinct bands of approximately 0.8 kb (Long) and 0.4 kb (Short). For sequencing, PCR fragments from 72 h samples were excised and purified using the FAST GENETM Gel/PCR Purification Kit (Nippon Genetics, F-91302).

Isoform-specific primers were designed: αGLT-Long primers targeted its unique sequence, and αGLT-Short primers recognized the Iα–IgA constant region splice junction unique to the short isoform. To validate primer specificity, αGLT-Long and αGLT-Short fragments were cloned into the pBluescript SK+ vector using the In-Fusion cloning kit (Takara). PCR using these plasmids as templates confirmed that the αGLT-Long primer set amplified only the Long plasmid, and the αGLT-Short primer set amplified only the Short plasmid.

Standard curves for absolute quantification were generated using plasmid DNA containing αGLT-Long or αGLT-Short, which was serially diluted from 100 pg (≈5 × 10⁷ molecules for a 3 kb dsDNA plasmid) in 10-fold steps. Isoform-specific real-time PCR was then performed, and the resulting curves enabled quantification of intracellular copy numbers of αGLT-Long and αGLT-Short transcripts. Primer sequences used for amplification and sequencing are listed in Table S1.

## Acknowledgements

We thank the members of the Departments of Biochemistry, Tohoku University Graduate School of Medicine for discussions and support and the Biomedical Research Core of Tohoku University Graduate School of Medicine for technical support.

## Author contributions

Conceptualization: Y.I., K.I.; Data curation: Y.I., H.S., K.F., D. S., K.I.; Formal analysis: Y.I., H.S., K.F., D. S., K.I.; Funding acquisition: Y.I., K.I; Investigation:; Y.I., H.S., K.F., D. S., K.I. Methodology: Y.I., H.S., K.F., D. S., K.I.; Resources: Y.I., K.I.; Supervision: K.I.; Visualization:; Y.I., K.I., Writing Y.I., K.I.

## Data and resource availability

All relevant data and details of resources can be found within the article.

---

**Table S1.**
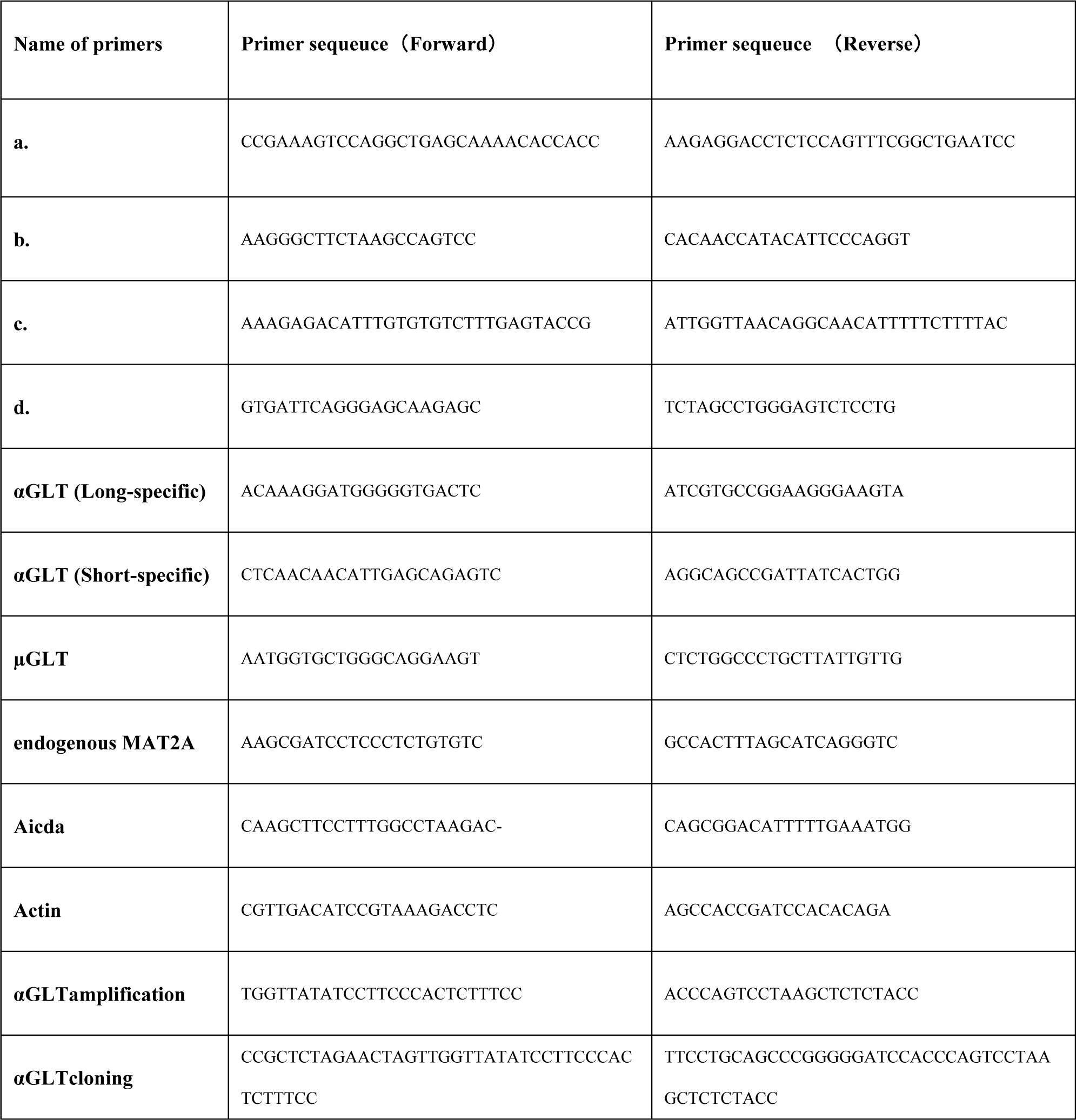
Primers used in this study.

